# Somatosensory Cortex of Macaque Monkeys is Designed for Opposable Thumb

**DOI:** 10.1101/2020.08.31.261115

**Authors:** Leslee Lazar, Prem Chand, Radhika Rajan, Hisham Mohammed, Neeraj Jain

## Abstract

The evolution of opposable thumb has enabled fine grasping ability and precision grip, which led to the capacity for fine manipulation of objects and refined tool use. Since tactile inputs to an opposable thumb are often spatially and temporally out of synch with inputs from the fingers, we hypothesized that inputs from the opposable thumb would be processed in an independent module in the primary somatosensory cortex (area 3b). Here we show that in area 3b of macaque monkeys, most neurons in the thumb representation do not respond to tactile stimulation of other digits and receive few intrinsic cortical inputs from other digits. However, neurons in the representations of other digits respond to touch on any of the four digits and are significantly more interconnected in the cortex. The thumb inputs are thus processed in an independent module, whereas there is significantly more inter-digital information exchange between the other digits. This cortical organization reflects behavioral use of the hand with an opposable thumb.

## Introduction

In many sensory areas of the brain, anatomically and functionally distinct inputs are processed in discrete modules. Examples include the ocular dominance columns in the primary visual cortex (V1) for inputs from the two eyes (Hubel and Wiesel 1962), segregation of trigeminal and spinal nerve inputs in the somatosensory area 3b, thalamus and medulla (Kaas et al. 2002), blobs in the supragranular layers of V1 for the color inputs (Horton 1984), and stripe-like modules in the second visual area (V2) (Roe and Tso 1997). Such modules constitute a network of dedicated neurons that process a subset of the sensory inputs, which code for either spatially segregated or sub-modality specific features. This helps reduce the dimensionality of the complex inputs for more efficient information processing. Outputs of these modules are then integrated with other independently processed inputs. Thus, information-processing modules provide an efficient means for analyzing the inputs, while helping the brain keep track of their nature and source.

The hand is the primary organ for object manipulation in primates (Macfarlane and Graziano 2009). Similar to humans, the opposable thumb of macaque monkeys gives them the ability to form a precision grip (Heffner and Masterton 1975; Macfarlane and Graziano 2009). The opposable thumb also provides an ability to grasp a large object with the thumb on one side and the fingers and palm on the other side of the object. In natural day-to-day behavior, humans and macaques use many variations of power grips, like, hand wrap, finger-splayed wrap and thumb-to-finger grip (Marzke 1997; Macfarlane and Graziano 2009). During these types of grips, the independently controlled opposable thumb and the fingers touch opposite surfaces of the objects, leading to a spatial and temporal discordance of the tactile inputs from the opposable thumb and the fingers. We, therefore, hypothesized that for efficient information processing, the evolution of opposable thumb (digit 1 or D1) would lead to the emergence of an independent information-processing thumb module in the somatosensory cortex. Conversely, there would be more information exchange between fingers, i.e. digit 2 to digit 5 (D2-D5), because they tend to be used together and are more likely to encounter similar inputs (Marzke 1997). We tested this hypothesis first by electrophysiologically determining the extent to which neurons in each digit representation in the primary somatosensory cortex, i.e. area 3b, are activated by tactile stimulation of other digits. In the second set of experiments, we used neuroanatomical techniques to establish connectional independence of the separate ‘thumb module’. Previous studies suggest that thalamocortical projections between individual digit representations in the ventroposterior lateral (VPL) nucleus of the thalamus and those in area 3b show a high degree of fidelity, i.e. there is very little cross-projections between the individual digits representations in the thalamus and area 3b (Jones et al. 1982). Therefore, we compared intrinsic anatomical connectivity between the thumb and other digit representations in area 3b.

## MATERIALS AND METHODS

Eight adult macaque monkeys (7 *Macaca mulatta* and 1 *Macaca radiata*; two males and six females, 4.4-7.3 kg) were used for these experiments (Table 1). Electrophysiological experiments were performed in four monkeys (07-68NM, NM56 08-56NM, 09-15NM, 10-31NM), and the intrinsic connections were determined in five monkeys (09-47NM, 09-51NM, 10-15NM, 10-31NM, 11-22NM). For monkey 10-31NM, both the hemispheres were used, left for the anatomical and right for the electrophysiological experiments. The animals were kept in a 12-hour dark and 12-hour light cycle. All animal protocols were approved by Institutional Animal Ethics Committee of National Brain Research Centre, and Committee for the Purpose of Control and Supervision of Experiments on Animals (CPCSEA), Government of India.

**Table 1:**
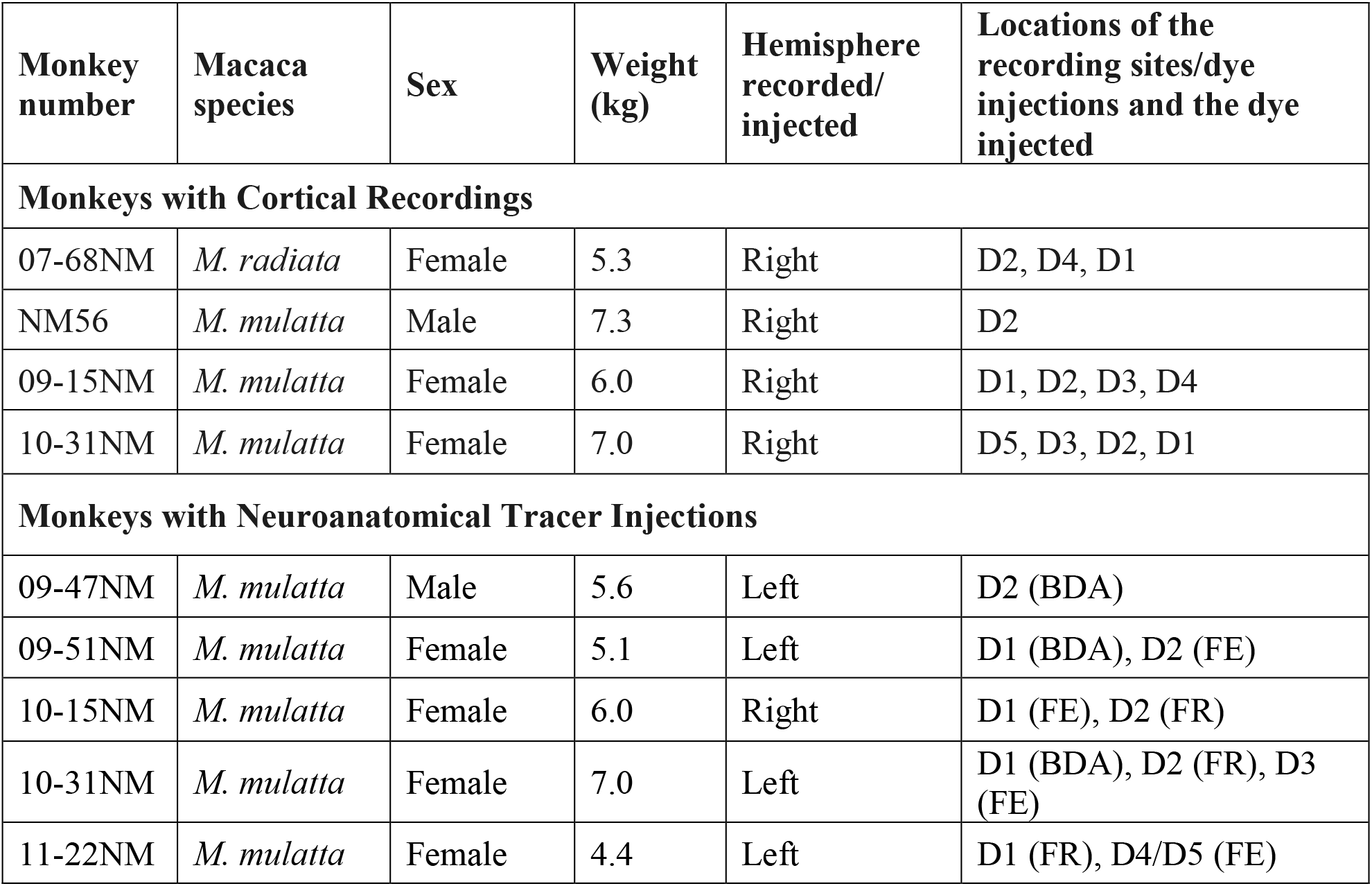
Details of monkeys in which electrophysiological recordings were performed or neuroanatomical tracers were injected in area 3b. The location of the injection site and the injected tracers are also shown, FR, fluororuby; FE, fluoroemerald and BDA, biotinylated dextran amine. Both the hemispheres of the monkey 10-31NM were used for different experiments.

| Monkey number | Macaca species | Sex | Weight (kg) | Hemisphere recorded/injected | Locations of the recording sites/dye injections and the dye injected |
| --- | --- | --- | --- | --- | --- |
| **Monkeys with Cortical Recordings** |  |  |  |  |  |
| 07-68NM | *M. radiata* | Female | 5.3 | Right | D2, D4, D1 |
| NM56 | *M. mulatta* | Male | 7.3 | Right | D2 |
| 09-15NM | *M. mulatta* | Female | 6.0 | Right | D1, D2, D3, D4 |
| 10-31NM | *M. mulatta* | Female | 7.0 | Right | D5, D3, D2, D1 |
| **Monkeys with Neuroanatomical Tracer Injections** |  |  |  |  |  |
| 09-47NM | *M. mulatta* | Male | 5.6 | Left | D2 (BDA) |
| 09-51NM | *M. mulatta* | Female | 5.1 | Left | D1 (BDA), D2 (FE) |
| 10-15NM | *M. mulatta* | Female | 6.0 | Right | D1 (FE), D2 (FR) |
| 10-31NM | *M. mulatta* | Female | 7.0 | Left | D1 (BDA), D2 (FR), D3 (FE) |
| 11-22NM | *M. mulatta* | Female | 4.4 | Left | D1 (FR), D4/D5 (FE) |

### Multi-unit mapping

Detailed somatotopy of the hand representation in area 3b was determined prior to quantitative neuronal recordings and injections of the neuroanatomical tracers. Methods for mapping have been described in detail before (Tandon et al. 2009; Chand and Jain 2015). Briefly, the animals were anaesthetized to surgical levels with a mixture of ketamine (8 mg/kg, IM) and xylazine (0.4 mg/kg, IM). The head was fixed in a stereotaxic apparatus, a midline incision was made in the skin, and the underlying muscles were retracted. A craniotomy was performed over the central sulcus, and the dura was carefully incised and retracted to expose the cortex. A tungsten microelectrode (1MΩ at 1 kHz; Microprobe, Gaithersburg, USA) was inserted in the post-central gyrus perpendicular to the brain surface along the depth of the central sulcus. Receptive fields of neurons were determined every 400 or 500 μm as the electrode was advanced using a hydraulic microdrive (David Kopf Instruments, Tujunga, CA, USA). Glabrous skin and hairs were stimulated with handheld wooden probes or brushes. Neuronal responses to manual movements of the limb and digits, as well as to joint manipulations were also determined (Tandon et al. 2009; Chand and Jain 2015). Minimal area of the skin that elicited a strong, consistent response to light tactile stimulation was defined as the receptive field of the neurons. The locations of the receptive fields were marked on photographs of the body surface. A series of electrode penetrations, approximately 500 μm apart, were made in the mediolateral direction along the central sulcus. The locations of the electrode insertion sites in the brain were marked on an enlarged photograph of the brain surface. The heart rate and core body temperature were monitored continuously throughout the procedure. The body temperature was maintained at 37 C with a warm-water blanket placed under the monkey. Physiological saline was administered intravenously, alternating with dextrose (5% in saline) every 4 hours.

### Multi-channel recordings

After determining somatotopy of the digit representations in area 3b, silicon multi-electrode neural probes (10 mm long, 15 or 50 μm thick; Neuronexus Inc, Ann Arbor, USA) with multiple recording points along its length were used to simultaneously record from multiple locations along the depth of the central sulcus. The probe had 16 iridium recording sites (impendence 1.02 – 1.66 MΩ, area 413 μm^2^) located 200 μm apart. The probe width was 240 μm at the proximal end, which gradually tapered to 33 μm at the tip.

The probe was lowered in the cortex using a hydraulic micromanipulator (David Kopf Instruments, Tujunga, CA, USA). Locations of the recording sites in the cortex were ascertained by mapping the receptive fields and optimized, if needed, for obtaining data from the maximum number of recording points by adjusting the depth or reinserting the probe at a different location. For quantitative recordings, the skin of the digits was stimulated using an electromechanical Chubbuck type stimulator (Cantek Metatron Associates, Inc., Canonsburg, USA). The stimulus was delivered at 1 Hz using a probe made of vertical camel hairbrush with a tip cut to a diameter of 1 mm. The tip of the brush contacted the skin for 5 ms with a maximum force of <0.007 gwt, such that there was no visible indentation of the skin when observed under high power of the surgical microscope. The ramp rate of movement of the probe for stimulus onset was ~80 μ/ms, and for stimulus offset it was ~80 μ/ms. To stabilize the digits, the hand of the monkeys was placed on a custom-made dental acrylic mould, with the glabrous surface of the palm facing upwards. Fingernails were glued to the mould with surgical cyanoacrylate adhesive.

Recordings were made while sequentially stimulating the hand at 9 different sites (Fig. 1A). In a single recording session, 300 stimuli were delivered at each of the stimulation sites. The stimulated sites were on the distal phalange of D1, and distal and proximal phalanges of all other digits. For D1, only the distal phalange was stimulated because it was hard to deliver isolated stimuli on the small proximal phalange of D1. In addition, skin on the foot and hairs on the chin were stimulated as control (Supplementary Figure 1). After stimulating all the sites, the multi-channel probe was inserted at a different location and the sequence of stimulations was repeated. In each monkey 3 or 4 penetrations were made.

**Figure 1.**
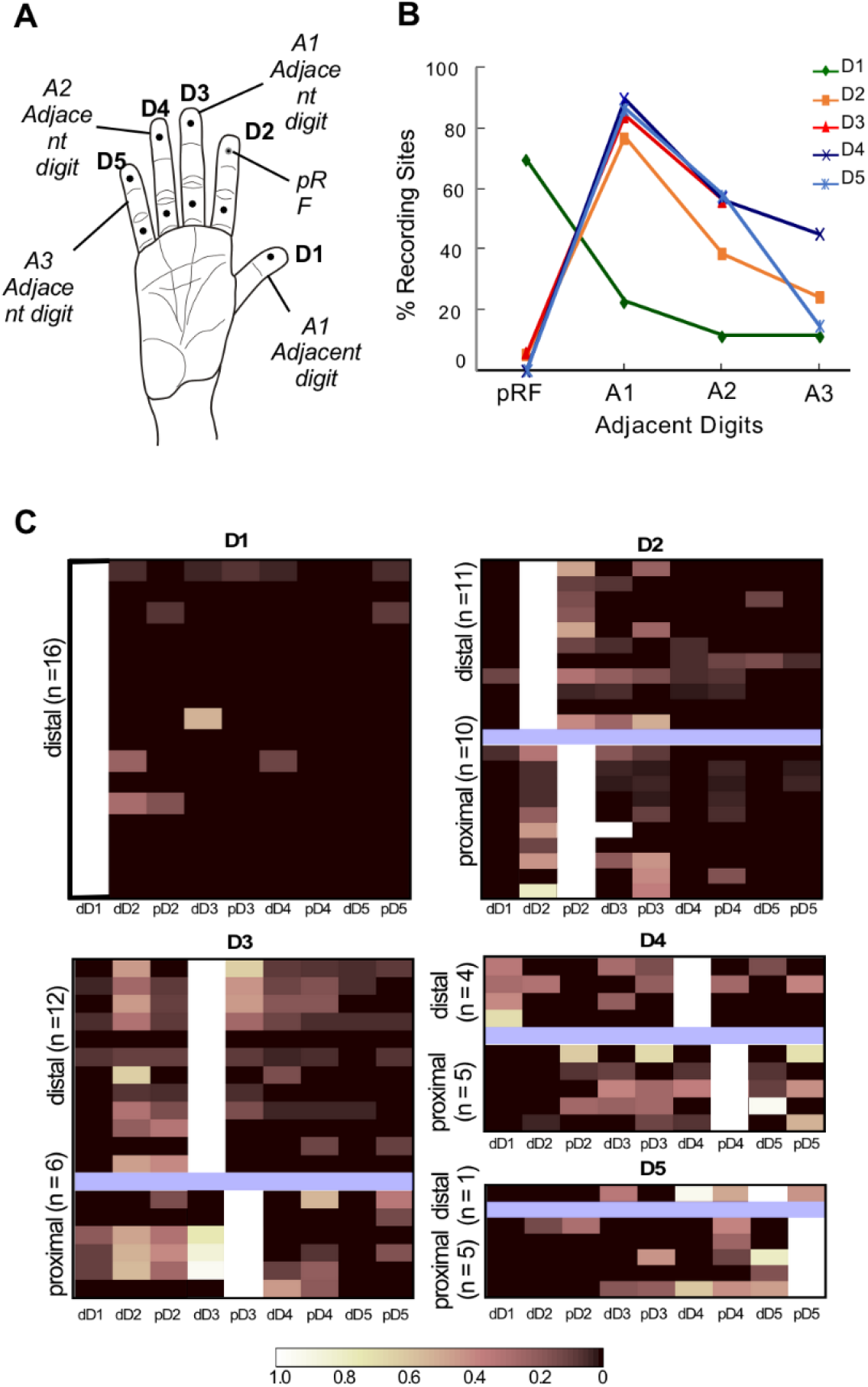
**(A)** An outline drawing of the glabrous surface of a hand of macaque monkey showing sites that were mechanically stimulated (dots). Digits are labeled in the black font. The nomenclature used for adjacency of digits with respect to the pRF (the circled dot on dD2 in this example) is shown in the italic font. **(B)** A graph showing the percentage of recording sites where neurons responded to touch only on pRF, and also on the A1 adjacent digit, the A2 adjacent digit, and the A3 adjacent digit. Data from A4 adjacent digit in not considered because neurons at only two recording sites had responses to touch on an A4 adjacent digit. **(C)** Matrix plots showing neuronal responses at different recording sites in different (D1 to D5) digit representations in area 3b (see label on top of each graph). The location of pRF on the distal or proximal phalange is shown on the y-axis. Each row in a plot shows responses of a single neuronal cluster to different stimulation sites as labeled on the x-axis. The color intensity represents response magnitudes normalized with respect to the response to stimulation of pRF (see the scale bar below). For digits, D2 - D5, the blue bar separates the recordings where pRF was on the distal (top) or the proximal (bottom) phalange.

Neuronal responses were amplified online, filtered by setting an appropriate threshold to remove low amplitude noise, digitized at 40 kHz and recorded using a multi-channel acquisition processor (MAP, Plexon Inc, USA). Spikes were detected using spike detection algorithm in the SortClient software (Plexon Inc, Dallas, USA) using a voltage-threshold trigger. Timestamps of the spikes and all the waveforms were saved on a computer for offline sorting and analysis.

After all the recordings were done, animals were euthanized with a lethal dose of pentobarbital (17.5 mg/kg, IV) and perfused transcardially with phosphate-buffered saline (0.9% NaCl in phosphate buffer, 0.1 M; pH 7.4), followed by 2% paraformaldehyde, and then by 2-4% paraformaldehyde in 10% sucrose. The brain was removed and cryoprotected in 30% buffered sucrose overnight. A block of the cortical tissue in the region of the central sulcus was frozen and cut into 50 or 60 μm thick sections on a sliding microtome in an off-parasagittal plane that was perpendicular to the central sulcus (Tandon et al. 2009). Different series of sections were stained for cytochrome oxidase (CO) (Wong-Riley 1979), Nissl substance (Mohammed and Jain 2014), or acetylcholine esterase (AChE) activity (Geneser-Jensen and Blackstad 1971) to determine architectonic borders of area 3b and boundaries of the cortical layers. The brain sections were used to trace the path of the electrode tracks and to determine locations of the recording sites with respect to the cortical layers and areal boundaries (Fig. 2). Only those recording sites that were deemed to be in the middle layers of the cortex were considered for further analysis.

**Figure 2.**
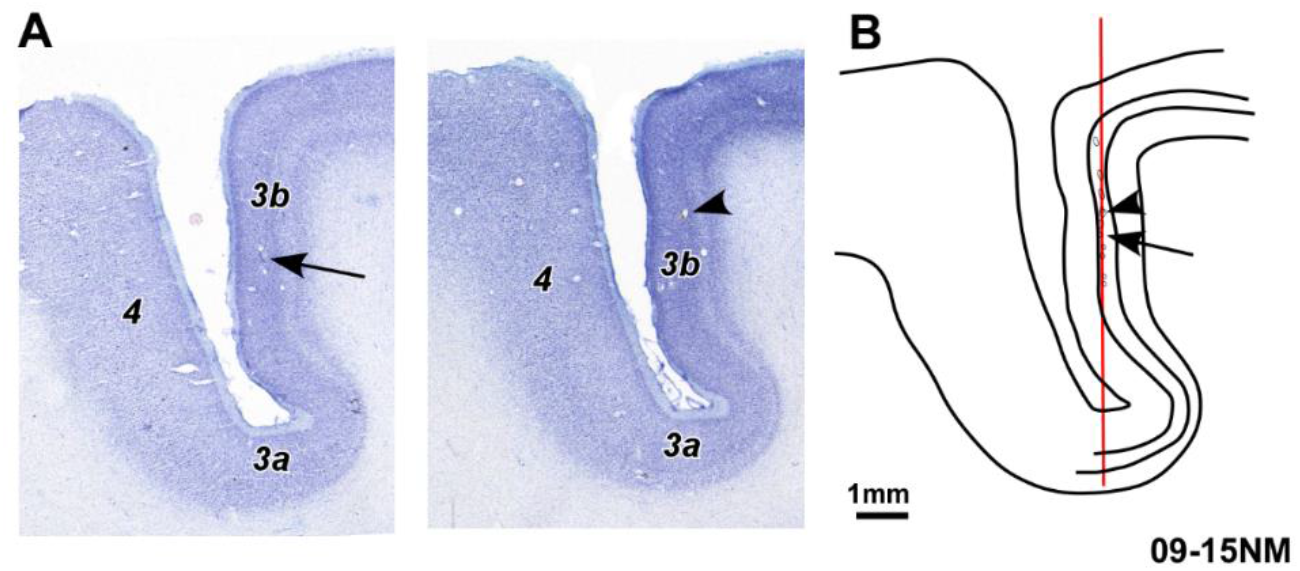
Reconstruction of a track of the multi-electrode probe. **(A)** A Nissl-stained section of the cortex in the region of the central sulcus cut in an off-parasagittal plane perpendicular to the central sulcus of monkey 10-31NM. Arrow points to part of the electrode track that was visible in this section. Locations of areas 3b, 3a, and 4 are marked. **(B)** An outline drawing of the section shown in ‘A’ with tracks from two adjacent sections overlaid (arrowheads) for the reconstruction of the entire track. Boundaries of layer 4 are marked. The grey line shows the path of the electrode as determined from these and additional sections. Scale bar, 1 mm.

#### Data analysis

The noise was removed from the recorded neuronal activity by first setting an appropriate threshold. Power spectral density graphs were examined to determine if there was any 50 Hz noise or other recording artefacts. If the noise was excessive, the data was discarded. The spikes were analyzed offline as multi-unit clusters (Offline Sorter; Plexon Inc., Dallas, USA) using principal component analysis (PCA) and verified by visual inspection (Nicolelis et al. 2003). Peri-stimulus time histograms (PSTH) were constructed with a bin size of 1 ms using Neuroexplorer software (Nex Technologies Inc., Madison, USA). The PSTH were smoothed using a boxcar filter of a width of 3 bins. Data was exported to MATLAB 2013A (The Mathworks Inc, Natick, USA) for further analysis. For quantification of neuronal response and other analyses, custom codes written in MATLAB, Microsoft Excel (Microsoft Inc., Washington, USA) and GraphPad Prism (GraphPad Software, Inc, California, USA) were used. Background firing rates of neurons were calculated as the mean firing rates during a 500 ms window before the stimulus onset. The response was analyzed in the post-stimulus time window of 100 ms. An excitatory response was considered to be present if the firing rate exceeded the mean background activity + 3 SD (standard deviation) in at least three consecutive bins. Peak response was defined as the firing rate of the bin with maximum value in the post-stimulus time window. The peak receptive field (pRF) was defined as a minimum area of the skin that elicited a strong, consistent response to light tactile stimulation in the classical mapping studies. Normalized peak firing rates for stimulations at different locations on the hand were obtained by dividing the peak firing rate for stimulation at that site by the peak firing rate for stimulation at the principal receptive field (pRF).

### Injections of neuroanatomical tracers

After determining somatotopy in area 3b by multi-unit mapping techniques described above, neuroanatomical tracers were injected at selected sites in the digit representations of five monkeys under sterile conditions (Table 1). The tracers were injected in the representations of D1, D2, D3, or D4/D5 at locations with robust neuronal responses (Chand and Jain 2015). Since the border between D4 and D5 is not always distinct in the electrophysiological maps, injections in D4 and D5 representations were designated as D4/D5 injections. Efforts were made to locate the injection sites in the centers of the digit representations with minimal contamination of the superficial cortex as the needle was advanced into the posterior bank of the central sulcus. At different sites, fluororuby (FR, 10% in saline; Molecular Probes, Invitrogen), fluoroemerald (FE, 10% FE in saline; Molecular Probes, Invitrogen), or biotinylated dextranamine (BDA, a mixture of 10% 10,000 MW BDA and 3% 3,000 MW BDA in 0.01 M sodium phosphate buffer; Molecular Probes, Invitrogen) was injected. FR and FE (0.03-0.08 μl) were pressure injected using a glass micropipette attached to a Hamilton syringe (Mohammed and Jain 2016). Injections of BDA were made iontophoretically (5 μA positive current, 7s on and 7s off pulses, total time 15 min). After injections, the brain was covered with sterile contact lenses (Acuvue, Johnson and Johnson, India), and the dura was folded back. A piece of sterile gel foam was placed on top of the dura and the craniotomy was sealed with a cap made from dental cement. The muscle and the skin were sutured in place. The animals were monitored closely during the recovery period and given enrofloxacin (5 mg/kg; IM), dexamethasone (2 mg/kg) and diclofenac (1.6 mg/kg) postoperatively for five days.

#### Histology

After 6-8 days the monkeys were euthanized and perfused with 2% paraformaldehyde as described above. In three of the monkeys (09-47NM, 10-15NM and 10-31NM), a detailed somatotopic map was obtained before perfusion, using techniques described above. After the brain was removed from the skull, the cortex was removed from the underlying tissue. A block of the cortex in the region of the central sulcus was manually flattened between glass slides, photographed and cryoprotected overnight in 30% sucrose. The flattened cortex was frozen and cut parallel to the pial surface on a sliding microtome into 40 μm thick sections. A series of alternate sections were either mounted unstained to visualize the fluorescently labeled neurons or processed for BDA (Reiner et al. 2000; Chand and Jain 2015). A second series of sections were stained for myelin using a modified Gallyas procedure (Jain et al. 1998) to visualize area 3b, the hand-face border, and the inter-digit borders (Chand and Jain 2015).

#### Data analysis

Locations of the labeled neurons were plotted using Neurolucida software (Microbrightfield, Williston, USA) with direct visualization under the microscope as described before (Mohammed and Jain 2014; Chand and Jain 2015). Outlines of the sections and locations of the blood vessels and other tissue artifacts were plotted to align the sections with each other. Adjacent sections stained for myelin were drawn at the same magnification, and myeloarchitectonic boundaries such as the hand-face border and the borders between digits representations, where visible, were marked (Jain et al. 1998; Chand and Jain 2015). In order to ascertain the locations of labeled neurons, plots of the labeled neurons and drawings from the myelin - stained sections were aligned using Canvas software (ACDSee Systems Inc., Seattle, USA). Only neurons determined to be in the hand representation of area 3b were considered for quantitative analysis (Chand and Jain 2015).

Finally, the electrophysiological map was overlaid on the drawing showing cell plots using the Canvas software. The number of labeled neurons in the representation of each digit in the hand representation of area 3b was counted. The mediolateral spread of the retrogradely labeled neurons within the hand region of area 3b was measured from the medial and lateral boundaries of the injection zone. Data was analyzed using Sigma Plot 13 (Systat Software, Inc., San Jose, USA) and plotted using GraphPad Prism (GraphPad Software Inc., CA, USA).

Retrogradely labeled neurons in the thalamic sections were also plotted in order to confirm whether the injection sites were located in the desired digit representation in area 3b by comparing with the previously published somatotopy of the hand in the ventroposterior nucleus (Qi et al. 2011), and to judge the adequacy of transport.

### Statistical Analysis

Statistical significance was calculated using Sigma Plot 13 (Systat Software, Inc., San Jose, USA). To statistically compare the cell numbers and spread of labeled neurons between the two groups, unpaired *t*-test was performed. Prior to this, normal distribution was ascertained using Shapiro-Wilk test.

Fisher’s exact test using a 2×2 contingency table was used to determine if there was a statistically significant difference between the number of recording sites responding to pRF alone for touch on D1 vs. rest of the digits, i.e. D2-D5.

For comparing the response magnitude and latency of neuronal responses to touch on pRF and other locations on the digits (A1, A2 and A3) statistical significance was tested using one-way Kruskal-Wallis test on ranks. Prior to this, a test for normality was conducted using Shapiro-Wilk test. If there was a significant difference among the groups, pair-wise analysis between each group was performed using Dunn’s Test.

## RESULTS

We first present our results on electrophysiological responses of neurons in different digit representations in area 3b to tactile stimulation on different digits. We then describe results on intrinsic connectivity between different digits representations in area 3b.

### Neuronal responses to stimulation at different locations on the hand

We recorded responses of neurons in the cortical representations of different digits in area 3b while the skin on each of the digits was subjected to periodic punctate stimuli (Fig. 1A). Simultaneous recordings from neuronal clusters at 16 different depths in 12 penetrations in the posterior bank of central sulcus were done from 4 anesthetized macaque monkeys using a Neuronexus multi-channel neural probe (see Methods; Table 1). Immediately before these quantitative recordings, area 3b was mapped using tungsten microelectrodes in order to delineate somatotopy of the hand representation in area 3b (Tandon et al. 2009) (see Methods).

Guided by the somatotopy of area 3b, the multi-channel probe was lowered into the desired location in the hand representation (Fig. 2), and the receptive field for each of the responsive channels was determined. The region of the skin that elicited the strongest response was designated as the primary receptive field (pRF). Simultaneous quantitative neural recordings from all the channels were then obtained while sequentially stimulating 9 different sites on the skin of the hand at 1 Hz using the tip of a camel-hair brush attached to an electromechanical Chubbuck-type stimulator (Kambi et al. 2014). The tip of the brush was cut flat to 1 mm diameter. The nine different locations on the glabrous skin of the digits were on the distal phalange of digit 1 (dD1), and distal (d) and proximal (p) phalanges of D2-D5 (Fig. 1). The tip of the brush touched the skin for 5 ms. The intensity of the stimulus was adjusted such that the tip of the brush touched the skin without any visible indentation when observed under a surgical microscope. The use of the brush helped ensure that deep stimuli were avoided because the hairs would visibly bend if the intensity of the touch were high.

We report here multi-unit activity recorded from 70 different recording sites in area 3b (Table 2). At the remaining sites, no neuronal responses were present, presumably because of the poor location of the recording sites in the cortex. Data was recorded using Multi-Neuron Acquisition Processor (MNAP) and analyzed offline (see Materials and Methods) (Kambi et al. 2014).

**Table 2.** The number of sites at which recordings were done for each of the primary receptive field (pRF).

| **pRF** | **Number of**<br>**recorded**<br>**sites** |
| --- | --- |
| dD1 | 16 |
| dD2 | 11 |
| pD2 | 10 |
| dD3 | 12 |
| pD3 | 6 |
| dD4 | 4 |
| pD4 | 5 |
| dD5 | 1 |
| pD5 | 5 |
| **TOTAL** | 70 |

Sixteen recordings were made in the D1 representation (Table 2). At the majority (68.8%, n = 11) of the sites, neurons responded to touch only on D1, i.e. the pRF (Fig. 1 B, C, 3). For these neurons, stimulation at any location on the hand other than D1 did not evoke any response. However, at five recording sites, neurons also responded to touch on other digits; neurons at 2 sites to touch on one additional digit, at 2 sites to two additional digits, and at 1 site to four additional digits (Fig. 1B).

**Figure 3.**
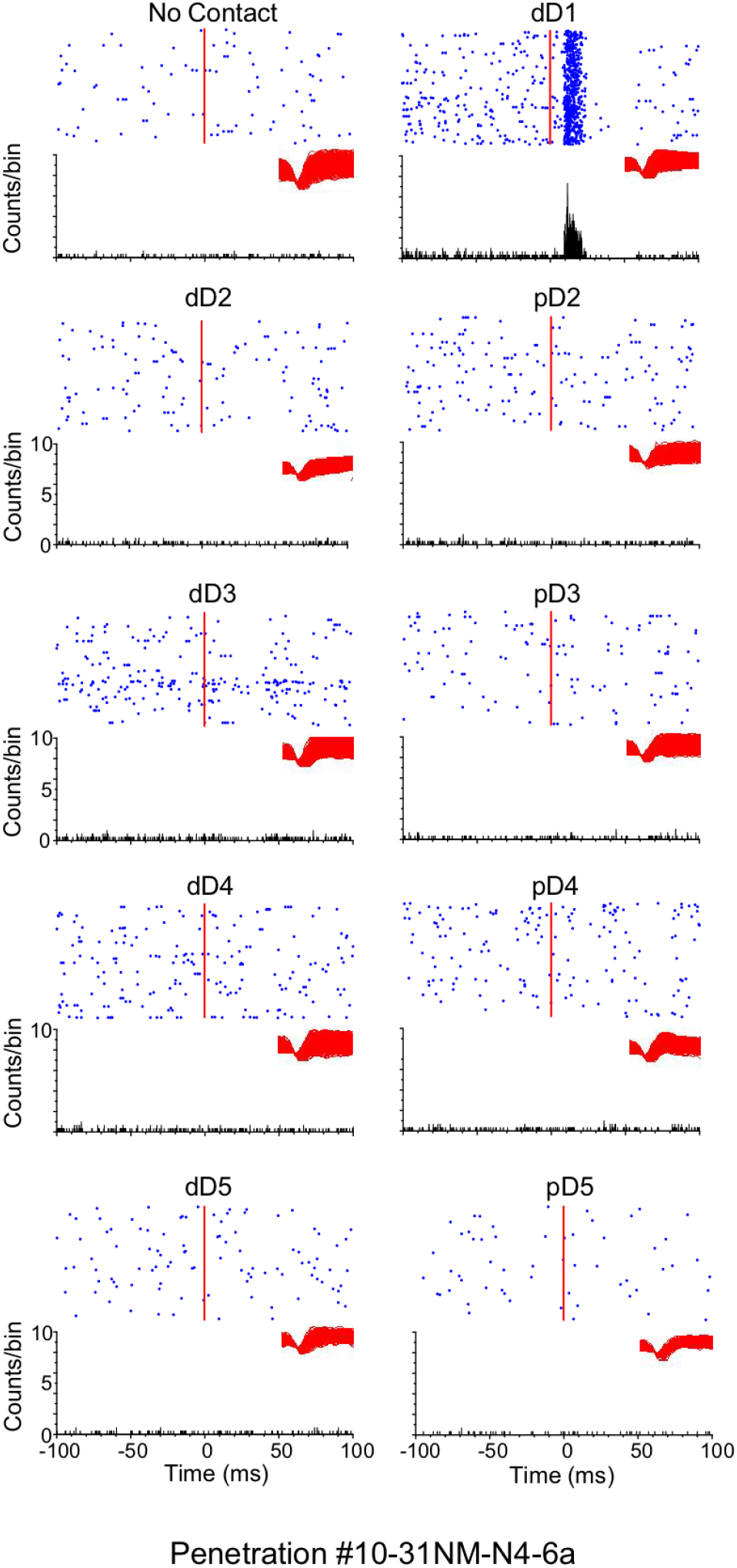
The response of neurons at the recording site ‘6a’ in penetration N4 of monkey 10-31NM. Each panel shows neuronal response (raster plots on the top and peri-stimulus time histograms (PSTH), below) to tactile stimulation at locations, which are labeled on top of each panel. The top left panel shows data when the stimulator was active but not touching the monkey, i.e. the spontaneous firing. In all panels, the data covers the duration from 100 ms before to 100 ms after the stimulus onset (vertical line, and zero on the x-axis). For PSTH, the y-axis shows the response magnitude (number of spikes in 1 ms bins, smoothed using a boxcar filter with a width of 3 bins). Each black dot in the raster plots denotes a single spike and the inset in black color shows the waveforms. For details, see “Methods’. At this recording location, touch only to D1 elicits a response.

In the representations of other digits, i.e. D2-D5, neuronal activity from a total of 54 sites was recorded (Table 2). Out of these, at 52 sites neurons responded to touch on multiple digits. This was significantly different as compared to D1 (p<0.0001; Fisher’s exact test). Only at two sites (i.e. 3.63%) responses to stimulation on pRF alone were observed, which were on dD2 and dD3. Neurons at 12 sites (22.2%) in D2-D5 representations also responded to touch on D1(Fig. 1B, C). Thus, most of the neurons in the representations of D2-D5 respond to touch on multiple digits, unlike those in D1 representation.

We also determined response magnitudes and the peak response latencies of neurons to touch on the skin of different digits (Fig. 4). Data showed that peak latencies of neurons for touch on pRF (n= 70) were significantly less as compared to touch on the adjacent digits (n=100) (Fig. 4A; p = <0.0001; Kruskal-Wallis test on ranks, see Materials and Methods). Similarly, the response magnitudes for touch on pRF were significantly higher as compared to touch on the adjacent digits (Fig. 4B; p = <0.0001; Kruskal-Wallis test on ranks). These results suggest that direct projections of thalamocortical inputs to multiple digit representations in area 3b are unlikely to mediate neuronal responses to non-pRF. If such overlapping thalamocortical inputs were mediating neuronal responses to the adjacent digits, these response parameters would likely be similar for touch on pRF and the adjacent digits. However, the presence of few short-latency responses for stimulation at locations other than pRF suggests that few such projections could be present.

**Figure 4.**
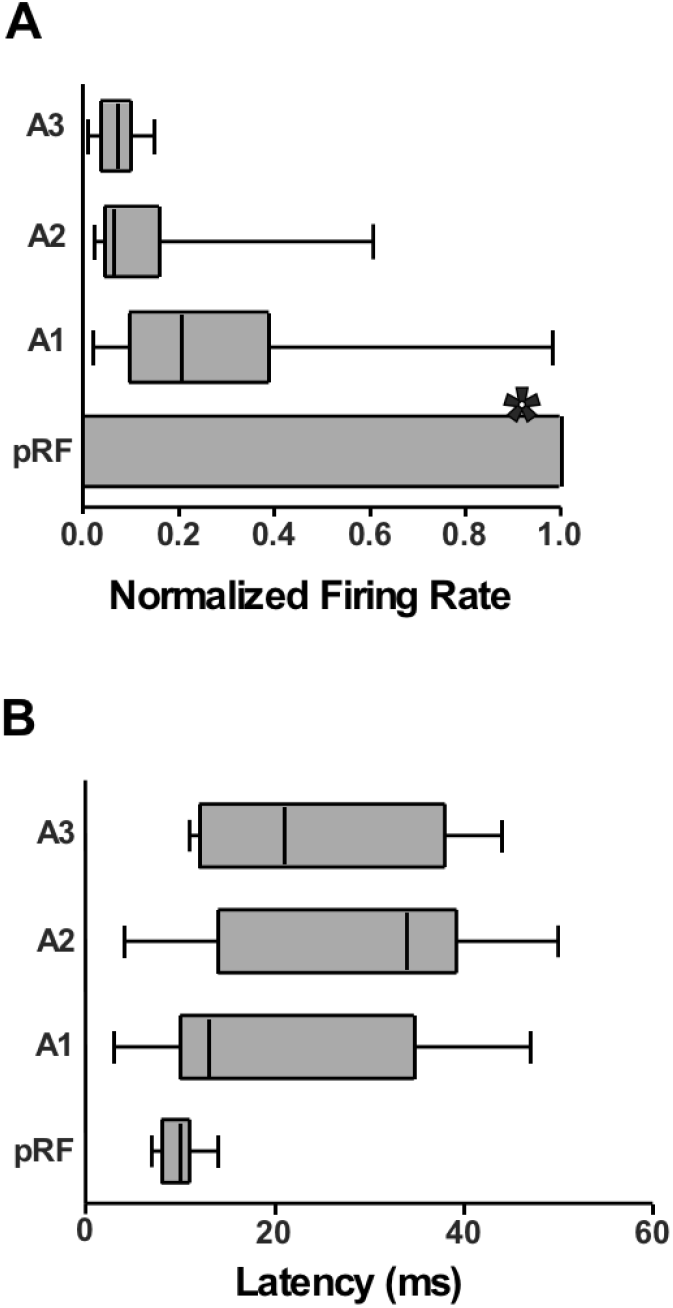
**(A)** Box plots showing response magnitudes at all the recording sites for tactile stimulation on pRF, A1, A2, and A3 adjacent digits normalized to the response magnitude to touch on pRF. The response magnitude to touch on pRF was significantly more as compared to touch on any of the adjacent digits (p<0.0001; one-way Kruskal-Wallis test of ranks). **(B)** Box plots showing peak response latencies for touch on pRF and the adjacent digits for all the recording sites. Peak latency to touch on pRF was significantly less as compared to touch on any of the adjacent digit (p<0.0001; One-way Kruskal-Wallis test of ranks). For both the box plots, the boxes enclose the 25^th^-75^th^ percentile, and the whiskers extend to the minimum and maximum values. The vertical line in the box marks the median value.

### Intrinsic connections of D1 representations are different as compared to other digits

We sought an anatomical basis for the presence of the D1 information-processing module. Intrinsic connections between different digit representations were determined by making small injections of neuroanatomical tracers - fluororuby (FR), fluoroemerald (FE) or biotinylated dextran amine (BDA) at ten different locations in different digit representations in 5 monkeys (Fig. 5, Table 1). In order to clearly delineate the extent of the tracer spread, cortex in the region of area 3b was flattened and cut into sections parallel to the pial surface (Jain et al. 2008; Chand and Jain 2015). One series of alternate sections was processed to visualize fluorescently labeled neurons or stained to determine the presence of the BDA label (Fig. 5). The other series of alternate sections were stained for myelin (Fig. 6; Jain et al. 1998), which we have previously shown to reveal anatomical borders between the digits, and between D1 and the face representations (Chand and Jain 2015). These borders are visible as myelin light septa. Using these myelin stained sections, we selected only those cases for analysis where injections sites were clearly confined to representation of a single digit for D1, D2 or D3. The border between D4 and D5 is usually not distinct in either electrophysiological maps or myelin-stained sections; therefore, the injection in D4 and D5 representations was designated as D4/D5 injection.

**Figure 5.**
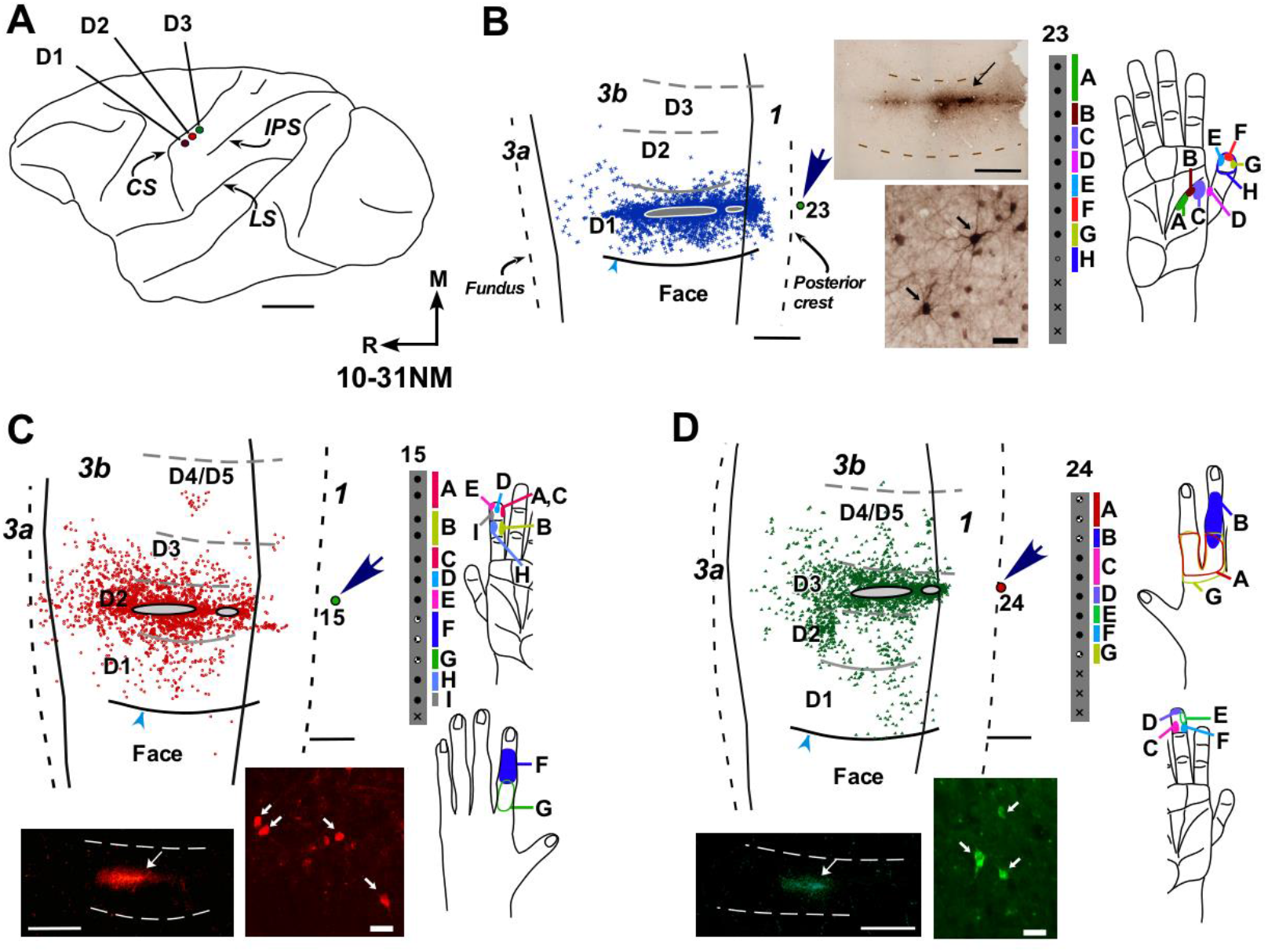
Intrinsic connections between different digit representations in area 3b. **(A)** An outline drawing of the lateral view of a macaque monkey brain showing locations of the injection sites (dots) in D1, D2 and D3 representations. Central sulcus (CS), intraparietal sulcus (IPS) and lateral sulcus (LS) are labeled for reference. **(B)** Location of the retrogradely labeled neurons following injection of BDA in the D1 representation. The white border with grey fill shows the core of the injection site, and each blue cross marks a labeled neuron. The dot labeled with number 23 (arrow) marks location of the recording site for which the receptive fields are shown in the far right figurine. Photomicrographs in the center show the injection site (above) and examples of the labeled neurons (below). On the far right, receptive fields of neurons recorded at an electrode penetration site close to the injection site marked with number 23. **(C and D)** Locations of labeled neurons following injections of FR in D2, and FE in D3 representations. Other conventions as for ‘B’ Scale bars, 1 cm for A,1 mm for B, C, D, and 20 μm for photomicrograph of the labeled neurons.

**Figure 6.**
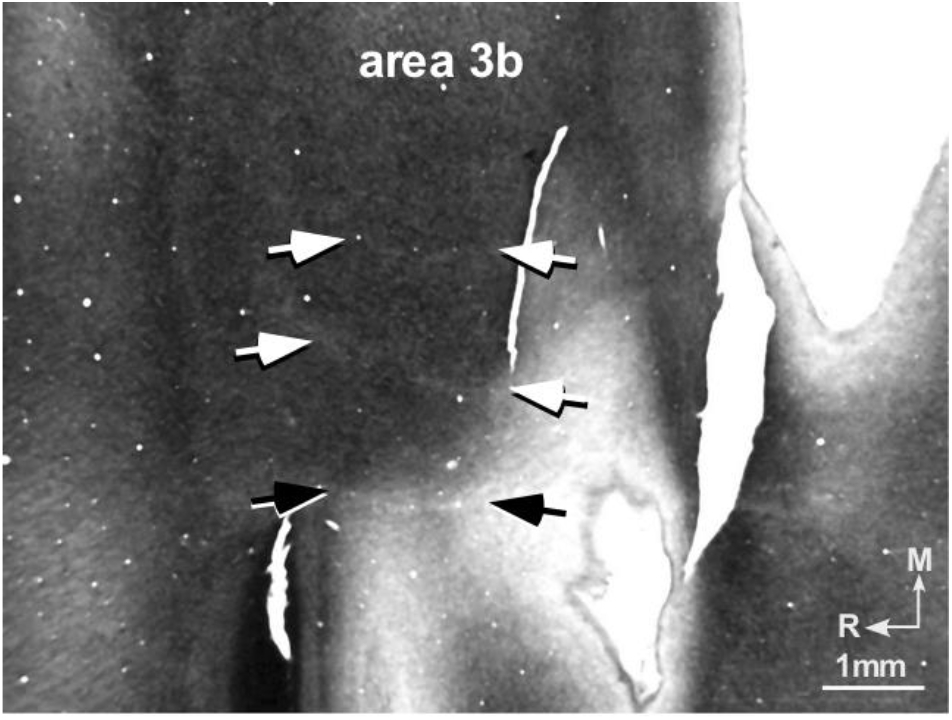
Photomicrograph of a myelin-stained section of the flattened cortex of macaque monkey 11-22NM. White arrowhead points to the myelin-light hand-face septum extending rostrocaudally, which separates medial hand and lateral face representation in area 3b. White arrow shows the fainter septa between digits D1–D2. Fundus and the posterior bank (dorsal lip) of the central sulcus are marked with dotted lines. M, medial; R, rostral.

The anatomical borders visible in myelin-stained sections helped us determine the precise locations of the retrogradely labeled neurons with respect to the digit representations. Our data show that for injections made in D1, the distribution of retrogradely labeled neurons is highly restricted and largely confined within the D1 representation (Fig. 5B). The labeled neurons were more widely distributed following injections in other digit representations (Fig. 5C-D). The average mediolateral spread of the labeled neurons in the hand representation for injections in D1 was 1.78±0.30 mm, for D2 it was 5.57±1.07 mm and for D3-D5, 5.28±0.62 mm (Fig. 7A). The difference in the mediolateral spread of labeled neurons between D1 and D2-D5 was significant (p < 0.0005, *t*-test). Thus neurons that project to the D1 representation are restricted to a narrow zone around the injection site, which was within D1 representation as confirmed by comparing with the anatomical borders revealed in the adjacent myelin stained sections (Fig. 5B).

**Figure 7.**
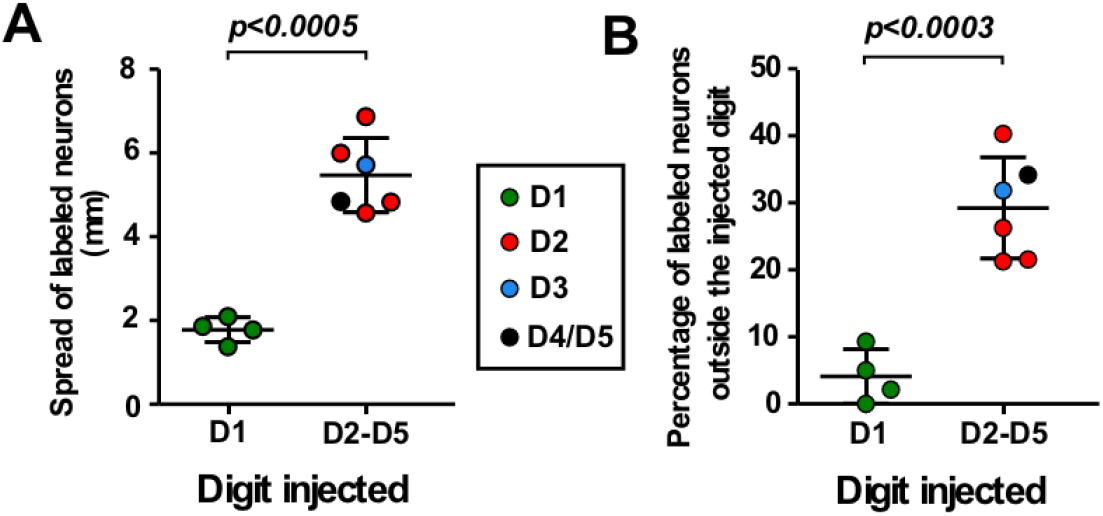
Distribution of neurons following injections of retrograde tracers in different digit representations. **(A)** Jittered scatter plots showing the maximum spread of labeled neurons following injection of the tracers in D1 (green dots) and digits D2-D5 (red, blue and black dots). Data for each digit is also shown separately coded by differently colored dots as per the key. Horizontal lines show mean (longer line) and the standard deviation. The spread of neurons is least for D1 and is significantly different from that for other digits. **(B)** Jittered scatter plots showing the percentage of total labeled neurons present outside the injected digit representation following injection of tracers in D1 (green dots) and digits D2-D5 (red, blue and black dots). Data for each digit is shown separately coded by differently colored dots as per the key. Longer horizontal lines show the mean and shorter, the standard deviation. For D1 injection, most of the neurons are present within D1 representation.

We also determined the number of labeled neurons within the injected digit representations as compared to the total number of the labeled neurons in the entire hand representation in area 3b. For injections in the D1 representation, only 4.1±4.0% of the labeled neurons were outside the D1 representation (Fig. 7B), the remaining 95.9±4.0% of the neurons were confined to the D1 representation. For the remaining digits, D2-D5, a significantly larger percentage of the labeled neurons was outside the injected digit representation (29.2±7.6%; p < 0.0004; t-test).

Thus, few neurons project to D1 from the other digit representations, suggesting a limited intrinsic exchange of information with neurons in other digit representations. Conversely, significantly more neurons project among representations of D2 – D5.

## DISCUSSION

We show that inputs from the digits to area 3b are processed in two functional modules – one related to inputs from the thumb (D1) and the other related to inputs from the remaining digits of the hand. We propose that this organization results in efficient information processing and reflects the use of the hand with an opposable thumb for object manipulation.

Earlier studies in awake macaque monkeys show that cortical neurons in D2-D5 (Lipton et al. 2010) and D2-D4 (Thakur et al. 2012) representations have multi-digit responses. Our results presented here corroborate these studies. However, in the previous studies responses of neurons in the D1 representation were not examined, which we show here to form an independent module, where neurons generally do not respond to stimulation on any other digit. Interestingly, an optical imaging study in macaque monkeys revealed that even in area 1, the D1 representation is segregated, whereas other digits show an overlapping representation, with a maximum overlap between digits 3-5 (Shoham and Grinvald 2001).

Subcortically, lemniscal inputs from different digits to VPL have minimal overlap (Jones 1983). Similarly, there is minimal overlap in the thalamocortical projections from different digit representations in VPL to area 3b (Padberg et al. 2009). Segregated digit specific modules for different digit representations can be discerned in VPL of the thalamus and the cuneate nucleus of the brain stem in suitable histological preparations (Jones 1983; Florence et al. 1989). In area 3b, although individual digit representations can be seen in the myelin stained section (Jain et al. 1998), our present results show that there are abundant intrinsic interconnections between D2-D5, which could mediate the multi-digit responses of neurons in these representations. Neurons in the D1 module get few inputs from neurons in D2-D5 representations.

New World monkeys such as owl monkeys and squirrel monkeys do not have an opposable thumb (Costello 1988). Available studies suggest that in New World monkeys, unlike macaque monkeys, neurons in the D1 representation are functionally and connectionally not different from the neurons in D2-D5 representations. In squirrel monkeys, Wang et al. (2013), report the presence of functional and anatomical connectivity between digits, including D1. Widespread integration of inputs between digit representations, including D1, has also been reported in owl monkeys (Reed et al. 2008; Reed et al. 2010). Injection of neuroanatomical tracer in the D1 representation of a squirrel monkey labeled neurons throughout the hand representation (Liao et al. 2013), similar to that for the other digits (Liao et al. 2013; Negyessy et al. 2013).

Developmentally, mapping studies in area 3b show that in New World monkeys, a detailed segregated representation of the digits is apparent soon after birth. However, in macaque monkeys, there are no clear borders between digit representation at birth (Krubitzer and Kaas 1988), which become apparent only by one month of age. We propose that early postnatal sensory experience likely reinforces the segregation of the thumb module in area 3b, similar to the development of segregated ocular dominance columns for the two eyes (Wiesel and Hubel 1965; LeVay et al. 1985). However, since tactile stimulation begins *in utero*, the overall somatotopy could be laid out even before birth (Jain et al. 1995; Dall’Orso et al. 2018).

Modular organization in other parts of the somatosensory system of monkeys has been reported. For example, inputs from the face, teeth, and tongue are processed in segregated modules in area 3b (Jain et al. 2001; Iyengar et al. 2007). A prominent myelin-light septum separating the trigeminal and spinal inputs can be seen in myelin stained sections of the flattened cortex (Jain et al. 1998). Moreover, there are very few intrinsic connections across the hand-face border (Manger et al. 1997; Fang et al. 2002; Chand and Jain 2015). Interestingly, the most prominent inter-digit myelin-light septum in the hand representation in area 3b is between D1 and D2 representations that have few connections crossing the border. Other digit representations that are separated by fainter myelin-light septa (Jain et al. 1998; Qi et al. 2011), form a more interconnected single module. Examination for the presence of myelin-light septa in macaque monkeys shows that these are present in 2-4 week old monkeys (Qi and Kaas 2004). Thus myelin-light septa separating digit representation do not always directly correlate with the physiological properties of neurons or even intrinsic connections.

Although the thumb representation is the largest of all the digits in area 3b of macaque monkeys, there are no differences in the density of Meissner’s corpuscles, tactile sensitivity, or receptive field sizes of neurons between the thumb and other digits (Chand and Jain 2015; Verendeev et al. 2015; Sathian and Zangaladze 1996; Sur et al. 1980). However, the opposable thumb has a unique role in behavior, which is vital for manual dexterity and success of a variety of grips (Napier 1956). Therefore, we propose that differences in the cortex between the thumb and the other digits in inter-digital intrinsic connectivity and neuronal response properties reflect the thumb opposability, i.e. the evolutionary change in the hand configuration and its use. This organization helps in efficient information processing of the two spatiotemporally disparate sets of inputs - one from the thumb and other from the remaining digits. The increased connectivity between the digits 2-5 would help integrate inputs from the co-activated peripheral receptors as these fingers are used together. Similar results have been found in the human primary motor cortex, where the structure of fMRI activations zones for individual digit representations follows the pattern of digit usage. The thumb shows a unique activation pattern, and the rest of the fingers show an overlapping pattern and similar dynamics during digit use (Ejaz et al. 2015).

It has been shown that sensory inputs and fine sensory discrimination abilities are critical for fine object manipulation and tactile skills such as precision grip. Monkeys with lesions of the dorsal columns of the spinal cord, which blocks ascending tactile inputs that enable fine discrimination, show a profound deficiency in the ability to perform precision grip (Glendinning et al. 1992; Qi et al. 2013). The movement representations for the digits in the primary motor cortex also change following lack of somatosensory inputs to the brain due to such injuries (Kambi et al. 2011). Comparisons can also be drawn with the organization of inputs from other sensory modalities in different mammalian species, where sensory inputs are processed in segregated functional modules. For example, in the primary visual cortex of monkeys, there are orientation columns, which are modules with similar orientation selectivity. Moreover, the orientation columns with similar orientation selectivity have a higher degree of horizontal connectivity, which helps integrate similar inputs from a wider area of the visual field (Weliky and Katz 1994; Weliky et al. 1995). Neurons in an orientation column tend to be influenced by stimuli in the visual fields of neighboring orientation columns (Dragoi et al. 2000). In the rat whisker barrel system, there is a higher degree of connectivity between the barrels corresponding to the same rostrocaudal row of whiskers as compared to the whiskers in the same column, i.e. those in the adjacent row (Hoeflinger et al. 1995). Electrophysiologically, stimulation of adjacent whiskers in the same row elicits a stronger response in neurons in a barrel corresponding to a whisker, as compared to when neighboring whiskers in the adjacent rows are stimulated, providing better functional connectivity (Rema and Ebner 1999).

Our results show that in area 3b of macaque monkeys, inputs from all the digits of the hand are not organized equivalently. Inputs from the opposable thumb are processed in a separate independent module, whereas neurons receiving inputs from the other four digits process the inputs in a more integrated manner. This organization is consistent with the behavioral use of a hand with an opposable thumb for grasping and manipulating objects, rather than any special tactile abilities of the thumb.

## Acknowledgements

This work was supported by the Department of Biotechnology, Government of India grant BT/PR7180/MED/30/907/2012, and funds from National Brain Research Centre to N.J.; P.C. is Research Associate, supported by Council of Scientific and Industrial Research, Government of India, grant 9/821(0034)/2011-EMR-I. We thank Dr. Niranjan Kambi for help with some of the electrophysiological and surgical procedures; Mr. Mithlesh Singh and Mr. Hari Shankar for histological and other support; and Dr. V. Rema for helpful comments on the manuscript.

## REFERENCES

Chand P, Jain N. 2015. Intracortical and Thalamocortical Connections of the Hand and Face Representations in Somatosensory Area 3b of Macaque Monkeys and Effects of Chronic Spinal Cord Injuries. The Journal of neuroscience : the official journal of the Society for Neuroscience. 35:13475–13486.

Costello MB, Fragaszy, D. M. 1988. Prehension in Cebus and Saimiri: 1. Grip Type and Hand Preference. American Journal of Primatology. 15:235–245.

Dall’Orso S, Steinweg J, Allievi AG, Edwards AD, Burdet E, Arichi T. 2018. Somatotopic Mapping of the Developing Sensorimotor Cortex in the Preterm Human Brain. Cerebral cortex. 28:2507–2515.

Dragoi V, Sharma J, Sur M. 2000. Adaptation-induced plasticity of orientation tuning in adult visual cortex. Neuron. 28:287–298.

Ejaz N, Hamada M, Diedrichsen J. 2015. Hand use predicts the structure of representations in sensorimotor cortex. Nature neuroscience. 18:1034–1040.

Fang PC, Jain N, Kaas JH. 2002. Few intrinsic connections cross the hand-face border of area 3b of New World monkeys. The Journal of comparative neurology. 454:310–319.

Florence SL, Wall JT, Kaas JH. 1989. Somatotopic organization of inputs from the hand to the spinal gray and cuneate nucleus of monkeys with observations on the cuneate nucleus of humans. The Journal of comparative neurology. 286:48–70.

Geneser-Jensen FA, Blackstad TW. 1971. Distribution of acetyl cholinesterase in the hippocampal region of the guinea pig. I. Entorhinal area, parasubiculum, and presubiculum. Z Zellforsch Mikrosk Anat. 114:460–481.

Glendinning DS, Cooper BY, Vierck CJ, Jr., Leonard CM. 1992. Altered precision grasping in stumptail macaques after fasciculus cuneatus lesions. Somatosensory & motor research. 9:61–73.

Heffner R, Masterton B. 1975. Variation in form of the pyramidal tract and its relationship to digital dexterity. Brain Behav Evol. 12:161–200.

Hoeflinger BF, Bennett-Clarke CA, Chiaia NL, Killackey HP, Rhoades RW. 1995. Patterning of local intracortical projections within the vibrissae representation of rat primary somatosensory cortex. The Journal of comparative neurology. 354:551–563.

Horton JC. 1984. Cytochrome oxidase patches: a new cytoarchitectonic feature of monkey visual cortex. Philos Trans R Soc Lond B Biol Sci. 304:199–253.

Hubel DH, Wiesel TN. 1962. Receptive fields, binocular interaction and functional architecture in the cat’s visual cortex. J Physiol. 160:106–154.

Iyengar S, Qi HX, Jain N, Kaas JH. 2007. Cortical and thalamic connections of the representations of the teeth and tongue in somatosensory cortex of new world monkeys. The Journal of comparative neurology. 501:95–120.

Jain N, Catania KC, Kaas JH. 1998. A histologically visible representation of the fingers and palm in primate area 3b and its immutability following long-term deafferentations. Cerebral cortex. 8:227–236.

Jain N, Florence SL, Kaas JH. 1995. GAP-43 expression in the medulla of macaque monkeys: changes during postnatal development and the effects of early median nerve repair. Brain research Developmental brain research. 90:24–34.

Jain N, Qi HX, Catania KC, Kaas JH. 2001. Anatomic correlates of the face and oral cavity representations in the somatosensory cortical area 3b of monkeys. The Journal of comparative neurology. 429:455–468.

Jain N, Qi HX, Collins CE, Kaas JH. 2008. Large-scale reorganization in the somatosensory cortex and thalamus after sensory loss in macaque monkeys. The Journal of neuroscience : the official journal of the Society for Neuroscience. 28:11042–11060.

Jones EG. 1983. Distribution patterns of individual medial lemniscal axons in the ventrobasal complex of the monkey thalamus. The Journal of comparative neurology. 215:1–16.

Jones EG, Friedman DP, Hendry SH. 1982. Thalamic basis of place- and modality-specific columns in monkey somatosensory cortex: a correlative anatomical and physiological study. Journal of neurophysiology. 48:545–568.

Kaas JH, Jain, N. Qi, H. 2002. The Organization of the Somatosensory System in Primates. In: The Somatosensory System. Deciphering the Brain’s own Body Image editor. Unites States of America: CRC Press.

Kambi N, Halder P, Rajan R, Arora V, Chand P, Arora M, Jain N. 2014. Large-scale reorganization of the somatosensory cortex following spinal cord injuries is due to brainstem plasticity. Nat Commun. 5:3602.

Kambi N, Tandon S, Mohammed H, Lazar L, Jain N. 2011. Reorganization of the primary motor cortex of adult macaque monkeys after sensory loss resulting from partial spinal cord injuries. The Journal of neuroscience : the official journal of the Society for Neuroscience. 31:3696–3707.

Krubitzer LA, Kaas JH. 1988. Responsiveness and somatotopic organization of anterior parietal field 3b and adjoining cortex in newborn and infant monkeys. Somatosensory & motor research. 6:179–205.

LeVay S, Connolly M, Houde J, Van Essen DC. 1985. The complete pattern of ocular dominance stripes in the striate cortex and visual field of the macaque monkey. The Journal of neuroscience : the official journal of the Society for Neuroscience. 5:486–501.

Liao CC, Gharbawie OA, Qi H, Kaas JH. 2013. Cortical connections to single digit representations in area 3b of somatosensory cortex in squirrel monkeys and prosimian galagos. The Journal of comparative neurology. 521:3768–3790.

Lipton ML, Liszewski MC, O’Connell MN, Mills A, Smiley JF, Branch CA, Isler JR, Schroeder CE. 2010. Interactions within the hand representation in primary somatosensory cortex of primates. The Journal of neuroscience : the official journal of the Society for Neuroscience. 30:15895–15903.

Macfarlane NB, Graziano MS. 2009. Diversity of grip in Macaca mulatta. Exp Brain Res. 197:255–268.

Manger PR, Woods TM, Munoz A, Jones EG. 1997. Hand/face border as a limiting boundary in the body representation in monkey somatosensory cortex. The Journal of neuroscience : the official journal of the Society for Neuroscience. 17:6338–6351.

Marzke MW. 1997. Precision grips, hand morphology, and tools. Am J Phys Anthropol. 102:91–110.

Mohammed H, Jain N. 2014. Two whisker motor areas in the rat cortex: evidence from thalamocortical connections. The Journal of comparative neurology. 522:528–545.

Mohammed H, Jain N. 2016. Ipsilateral cortical inputs to the rostral and caudal motor areas in rats. The Journal of comparative neurology. 524:3104–3123.

Napier JR. 1956. The prehensile movements of the human hand. The Journal of bone and joint surgery British volume. 38-B:902–913.

Negyessy L, Palfi E, Ashaber M, Palmer C, Jakli B, Friedman RM, Chen LM, Roe AW. 2013. Intrinsic horizontal connections process global tactile features in the primary somatosensory cortex: neuroanatomical evidence. The Journal of comparative neurology. 521:2798–2817.

Nicolelis MA, Dimitrov D, Carmena JM, Crist R, Lehew G, Kralik JD, Wise SP. 2003. Chronic, multisite, multielectrode recordings in macaque monkeys. Proc Natl Acad Sci U S A. 100:11041–11046.

Padberg J, Cerkevich C, Engle J, Rajan AT, Recanzone G, Kaas J, Krubitzer L. 2009. Thalamocortical connections of parietal somatosensory cortical fields in macaque monkeys are highly divergent and convergent. Cerebral cortex. 19:2038–2064.

Qi HX, Gharbawie OA, Wong P, Kaas JH. 2011. Cell-poor septa separate representations of digits in the ventroposterior nucleus of the thalamus in monkeys and prosimian galagos. The Journal of comparative neurology. 519:738–758.

Qi HX, Gharbawie OA, Wynne KW, Kaas JH. 2013. Impairment and recovery of hand use after unilateral section of the dorsal columns of the spinal cord in squirrel monkeys. Behavioural brain research. 252:363–376.

Qi HX, Kaas JH. 2004. Myelin stains reveal an anatomical framework for the representation of the digits in somatosensory area 3b of macaque monkeys. The Journal of comparative neurology. 477:172–187.

Reed JL, Pouget P, Qi HX, Zhou Z, Bernard MR, Burish MJ, Haitas J, Bonds AB, Kaas JH. 2008. Widespread spatial integration in primary somatosensory cortex. Proc Natl Acad Sci U S A. 105:10233–10237.

Reed JL, Qi HX, Zhou Z, Bernard MR, Burish MJ, Bonds AB, Kaas JH. 2010. Response properties of neurons in primary somatosensory cortex of owl monkeys reflect widespread spatiotemporal integration. Journal of neurophysiology. 103:2139–2157.

Reiner A, Veenman CL, Medina L, Jiao Y, Del Mar N, Honig MG. 2000. Pathway tracing using biotinylated dextran amines. J Neurosci Methods. 103:23–37.

Rema V, Ebner FF. 1999. Effect of enriched environment rearing on impairments in cortical excitability and plasticity after prenatal alcohol exposure. The Journal of neuroscience : the official journal of the Society for Neuroscience. 19:10993–11006.

Roe AW, Tso, D.Y. 1997. The functional architecture of area V2 in the macaque monkey: physiology, topography and connectivity. In: Cerebral Cortex Rockland etal, editor. New York Plenum Press.

Sathian K, Zangaladze A. 1996. Tactile spatial acuity at the human fingertip and lip: bilateral symmetry and interdigit variability. Neurology. 46:1464–1466.

Shoham D, Grinvald A. 2001. The cortical representation of the hand in macaque and human area S-I: high resolution optical imaging. The Journal of neuroscience : the official journal of the Society for Neuroscience. 21:6820–6835.

Sur M, Merzenich MM, Kaas JH. 1980. Magnification, receptive-field area, and “hypercolumn” size in areas 3b and 1 of somatosensory cortex in owl monkeys. Journal of neurophysiology. 44:295–311.

Tandon S, Kambi N, Lazar L, Mohammed H, Jain N. 2009. Large-scale expansion of the face representation in somatosensory areas of the lateral sulcus after spinal cord injuries in monkeys. The Journal of neuroscience : the official journal of the Society for Neuroscience. 29:12009–12019.

Thakur PH, Fitzgerald PJ, Hsiao SS. 2012. Second-order receptive fields reveal multidigit interactions in area 3b of the macaque monkey. Journal of neurophysiology. 108:243–262.

Verendeev A, Thomas C, McFarlin SC, Hopkins WD, Phillips KA, Sherwood CC. 2015. Comparative analysis of Meissner’s corpuscles in the fingertips of primates. J Anat. 227:72–80.

Wang Z, Chen LM, Negyessy L, Friedman RM, Mishra A, Gore JC, Roe AW. 2013. The relationship of anatomical and functional connectivity to resting-state connectivity in primate somatosensory cortex. Neuron. 78:1116–1126.

Weliky M, Kandler K, Fitzpatrick D, Katz LC. 1995. Patterns of excitation and inhibition evoked by horizontal connections in visual cortex share a common relationship to orientation columns. Neuron. 15:541–552.

Weliky M, Katz LC. 1994. Functional mapping of horizontal connections in developing ferret visual cortex: experiments and modeling. The Journal of neuroscience : the official journal of the Society for Neuroscience. 14:7291–7305.

Wiesel TN, Hubel DH. 1965. Extent of recovery from the effects of visual deprivation in kittens. Journal of neurophysiology. 28:1060–1072.

Wong-Riley M. 1979. Changes in the visual system of monocularly sutured or enucleated cats demonstrable with cytochrome oxidase histochemistry. Brain Res. 171:11–28.

