## SupplementalFig1 for "Somatosensory Cortex of Macaque Monkeys is Designed for Opposable Thumb"

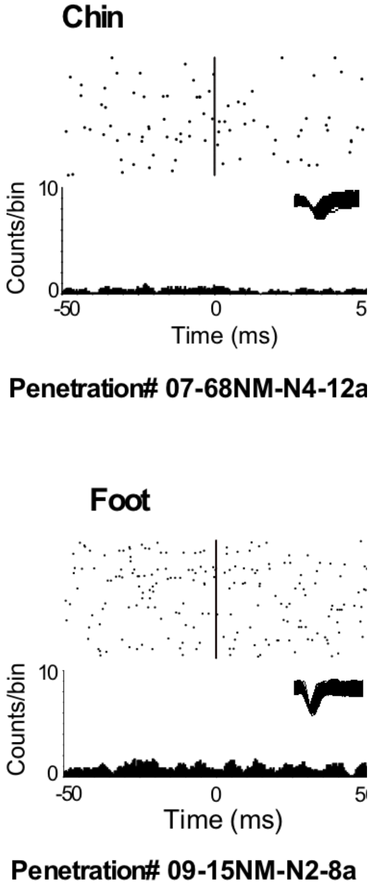


**Supplementary Figure S1.** Raster plots and PSTH for recording sites in the hand region of area 3b when hairs on the chin (top panel) or the skin of the foot (bottom panel) were stimulated as controls. No neuronal response was evoked.
